# Allocentric and egocentric cues constitute an internal reference frame for real-world visual search

**DOI:** 10.1101/2025.04.14.648618

**Authors:** Yan Chen, Zhe-Xin Xu

## Abstract

Visual search in natural environments involves numerous objects, each composed of countless features. Despite this complexity, our brain efficiently locates targets. Here, we propose that the brain combines multiple reference cues to form an internal reference frame that facilitates real-world visual search. Objects in natural scenes often appear in orientations perceived as upright, enabling quicker recognition. However, how object orientation influences real-world visual search remains unknown. Moreover, the contributions of different reference cues—egocentric, visual context, and gravitational— are not well understood. To answer these questions, we designed a visual search task in virtual reality. Our results revealed an orientation effect independent of set size, suggesting reference frame transformation rather than object rotation. By rotating virtual scenes and participants in a flight simulator, we found that allocentric cues drastically altered search performance. These findings provide novel insights into the efficiency of real-world visual search and its connection to multimodal cognition.

**Significance:** A central question in the behavioral sciences concerns how people efficiently perceive natural environments. Visual search exemplifies this challenge. While research has elucidated the basic mechanisms, traditional theories struggle to explain the remarkable efficiency in real-world scenes. Here, we examine a fundamental property of natural scenes: reference frames. Real-world objects typically appear in consistent orientations, suggesting that orientation may guide search. Yet, the influence of object orientation on real-world search—and which reference cues (egocentric, visual context, or gravitational) determine that orientation—remains unknown. We developed a novel virtual-reality paradigm to address these questions. We demonstrated that humans combine multiple reference cues to form an internal reference frame that guides visual search, providing a novel account of the efficiency of real-world search.

## Introduction

Imagine navigating a busy mall looking for your friend amidst the crowd, or spotting a bottle of juice among stacks of products in a grocery store. These seemingly mundane moments illustrate the remarkable efficiency of visual search, where the nervous system sifts through countless colors, shapes, and motions to find a specific target. How does our brain manage to find that needle in a haystack? What cues guide visual search in natural environments?

While decades of research have uncovered the elements of visual search, such as parallel and serial search^1–5^, traditional theories are limited to simplified, controlled settings and fail to explain search performance involving natural scenes. For example, feature integration theory^1,2^ predicts that visual search in the real world should be agonizingly slow, as it requires a step-by-step integration of numerous features. In practice, however, our brain performs these tasks with surprising speed and accuracy^5,6^.

Recent studies have revealed several factors contributing to this striking efficiency of real-world visual search, including top-down attention, stimulus saliency, semantic context, prior knowledge, and environmental statistics^7–13^. These factors establish a priority map that guides attention to specific features or spatial locations^9,10^. Another key element of natural scenes involves objects frequently occurring in consistent orientations^14–16^, which may aid visual search.

The role of orientation has been extensively studied in various recognition and discrimination tasks, with evidence suggesting that upright stimuli are processed more quickly and accurately^14,16–25^. Notably, response time scales linearly as an object’s orientation increasingly deviates from its canonical orientation, or as the orientation disparity between two objects increases^25–27^. These findings suggest two possible processes that underlie orientation dependency: mental rotation of objects and reference frame transformation^25^. Object mental rotation refers to a mental imagery process that rotates an object to its canonical orientation; in contrast, the latter process involves transforming a coordinate system to align with the object^25,27^. Neural mechanisms that underlie both processes have been proposed, including rotating the neural population vector for mental rotation^28,29^ and gain modulation for coordinate transformation^30–32^. While object recognition is likely a necessary step for visual search, it is unknown how orientation influences visual search for real-world objects and which process is involved.

Moreover, our brain represents the world in multiple reference frames^33–49^. For example, when tilting your head to peer through an aisle in a grocery store, the bottle of juice might appear tilted in your eyes. Nevertheless, the brain correctly identifies the bottle by integrating various reference cues, including how objects are oriented relative to the body (egocentric), the surrounding environment (visual context), and gravity^41,43,46,47,50–53^. It has been hypothesized that the brain forms an internal reference frame by combining multiple reference cues for object perception^24,25^. To date, the role of an internal reference frame and the contributions of different reference cues in real-world visual search remain unknown.

To address this major knowledge gap, we design a real-world visual search task in virtual reality. We demonstrate a shorter response time (RT) and higher accuracy for upright objects than for those oriented horizontally. This orientation dependency is independent of set size, indicating a process outside of serial search is involved, namely the transformation of an internal reference frame rather than individual object rotation. By pairing a flight simulator with a head-mounted display, we independently manipulate the visual scene and participants’ body orientation, disentangling the contributions of different reference cues that are normally overlapped. We show that orientation dependency is significantly altered by visual context and gravitational cues, demonstrating the involvement of multisensory signals and high-level cognitive processes. This study provides a systematic examination of how object orientation influences visual search in naturalistic environments, offering insights into visual processing in the natural world.

## Results

### Orientation dependency in real-world visual search

We started by asking how object orientation influences the performance of real-world visual search. We designed a psychophysical task in a virtual reality system that consisted of a head-mounted display and a flight simulator (see Methods). Objects were presented in a natural scene in the virtual environment, and participants were asked to report whether a cued target was present (Figure 1). Eight experimental conditions were introduced, including the combinations between two set sizes (4 and 9), two object orientations (0°, upright; 90°, horizontal), and target presence (present or absent).

**Figure 1.**
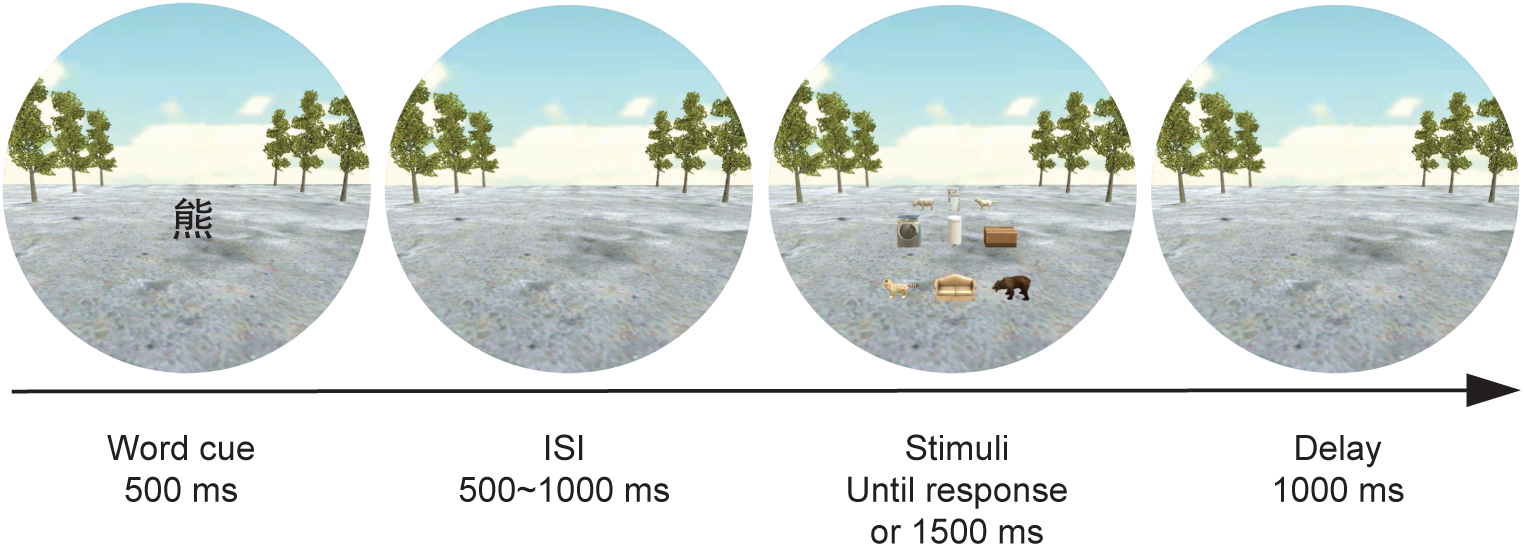
Procedure and stimuli. At the beginning of each trial, a word cue was presented at the center of the display (e.g., “Bear”), indicating the present trial’s search target. After a variable inter-stimulus interval (ISI), objects appeared for 1500 ms or until a response was made by the participant. The next trial started after a 1000-ms delay.

All participants achieved accuracies well above chance, ranging from 81.4% to 95.6%, with a mean of 89.4%. The hit rate ranged from 71.2% to 94.7%, with a mean of 84.4%, and the false alarm rate ranged from 3.2% to 10.1%, with a mean of 5.6%. Discriminability ranged from 1.89 to 3.43, with a mean of 2.65.

We used a linear mixed-effects model to quantify the effects of various factors on RT while accounting for variability across participants and object categories. The model included object orientation, target presence, set size, and interactions between them as fixed effects. Subject identity and object category were included as random effects (see Methods). We found a significant effect of object orientation on normalized RT, with longer RT for upright objects than for horizontal ones (Figure 2A; estimate=0.002, 95%CI=[0.00055, 0.0034], *t*(10029)=2.7, *p*=0.00695). Participants also showed higher accuracy for upright objects than for horizontal ones (Figure 2B; estimate=-5.02×10^−4^, 95%CI=[−9.69×10^−4^, −3.63×10^−5^], *t*(11146)=-2.11, *p*=0.0346), suggesting that the difference in RT between these two object orientations cannot be explained by a speed-accuracy trade-off.

**Figure 2.**
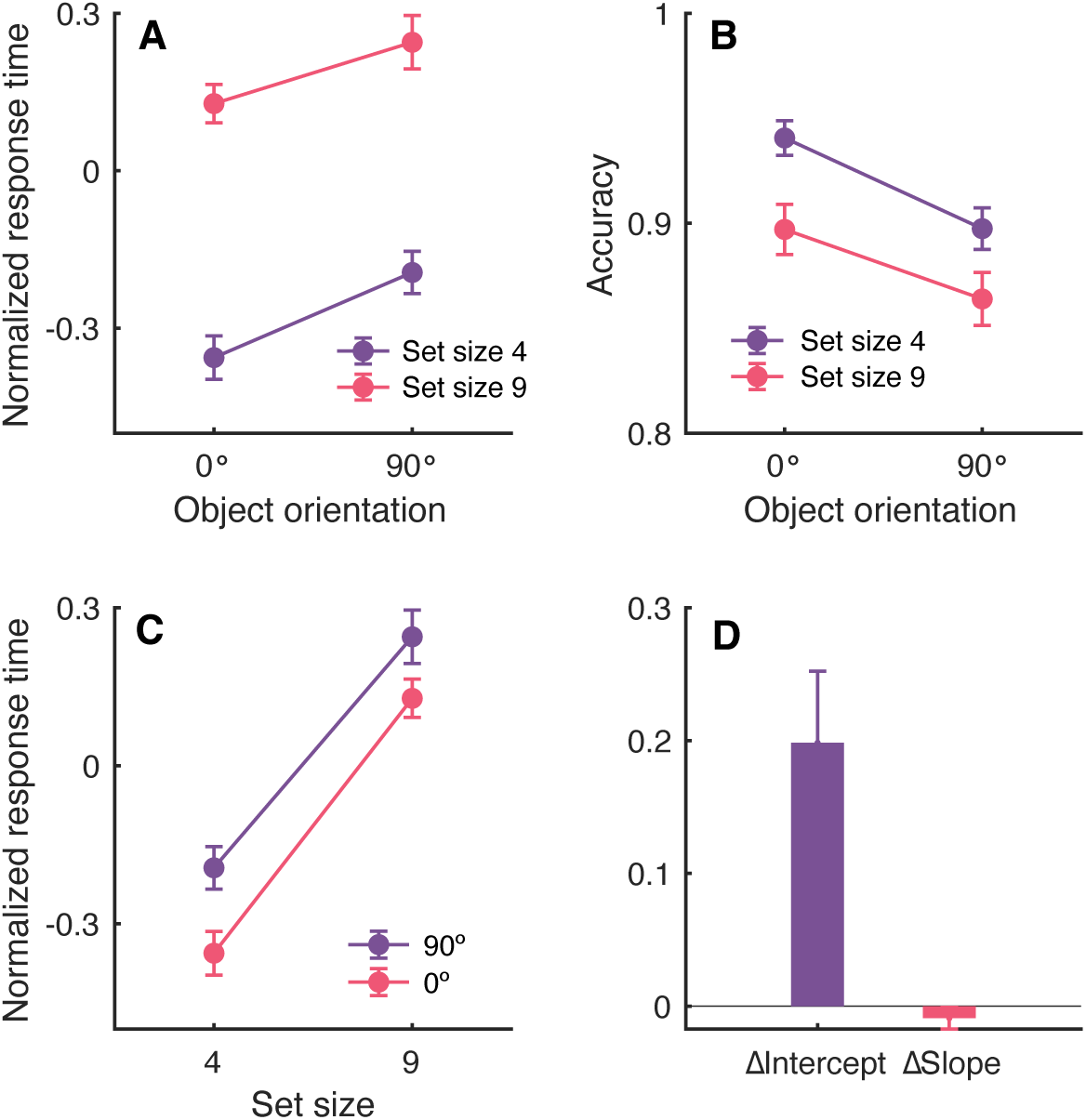
Results for Experiment 1. A, Normalized response time was longer for horizontally oriented objects (90°) than for upright objects (0°) for set sizes 4 (purple line) and 9 (red line). B, The accuracy was higher when objects were upright than when they were horizontal. Format as in A. C, Normalized response time increased as a function of set size for upright (red line) and tilted objects (purple line). D, Difference in intercept and slope of the response time–set size function (lines shown in C) between upright and horizontal objects. Error bars represent S.E.M.

These results demonstrate that real-world visual search is faster and more accurate for upright objects than for horizontal objects, suggesting an orientation dependency in this task. In addition, we found that normalized RT scaled with set size (Figure 2C; estimate=0.129, 95%CI=[0.116,0.142], *t*(10029)=19.1, *p*=1.26×10^−79^), which suggests that a serial search was involved.

### Internal reference frame facilitates visual search

What cognitive process might give rise to this orientation dependency in visual search? Previous studies on object recognition suggest two possibilities^25,27^: individual object rotation and internal reference frame transformation (Figure 3A). As discussed above, the effect of set size on RT indicates a serial search, in which individual objects are processed one at a time^2^. If the observer mentally rotates each object during this serial process^54^, we expect set size to have a larger effect on RT for tilted objects than for upright ones. In other words, each additional object adds to the total mental rotation time, resulting in a steeper slope of the RT–set size function for tilted objects (Figure 3A and B, bottom row). Note that any process that applies to individual objects necessarily predicts an orientation effect on the slope, not the intercept. For example, if orientation acts as a gain on object processing, orientation dependency should also increase with set size because of the additional process time during serial search. Alternatively, the observer may rotate an internal reference frame to align with the objects and maintain its orientation throughout the serial process^25^. Because reference frame transformation happens before serial search and would be constant regardless of the number of objects present, we expect the effect of object orientation to be independent of set size (Figure 3A and B, top row).

**Figure 3.**
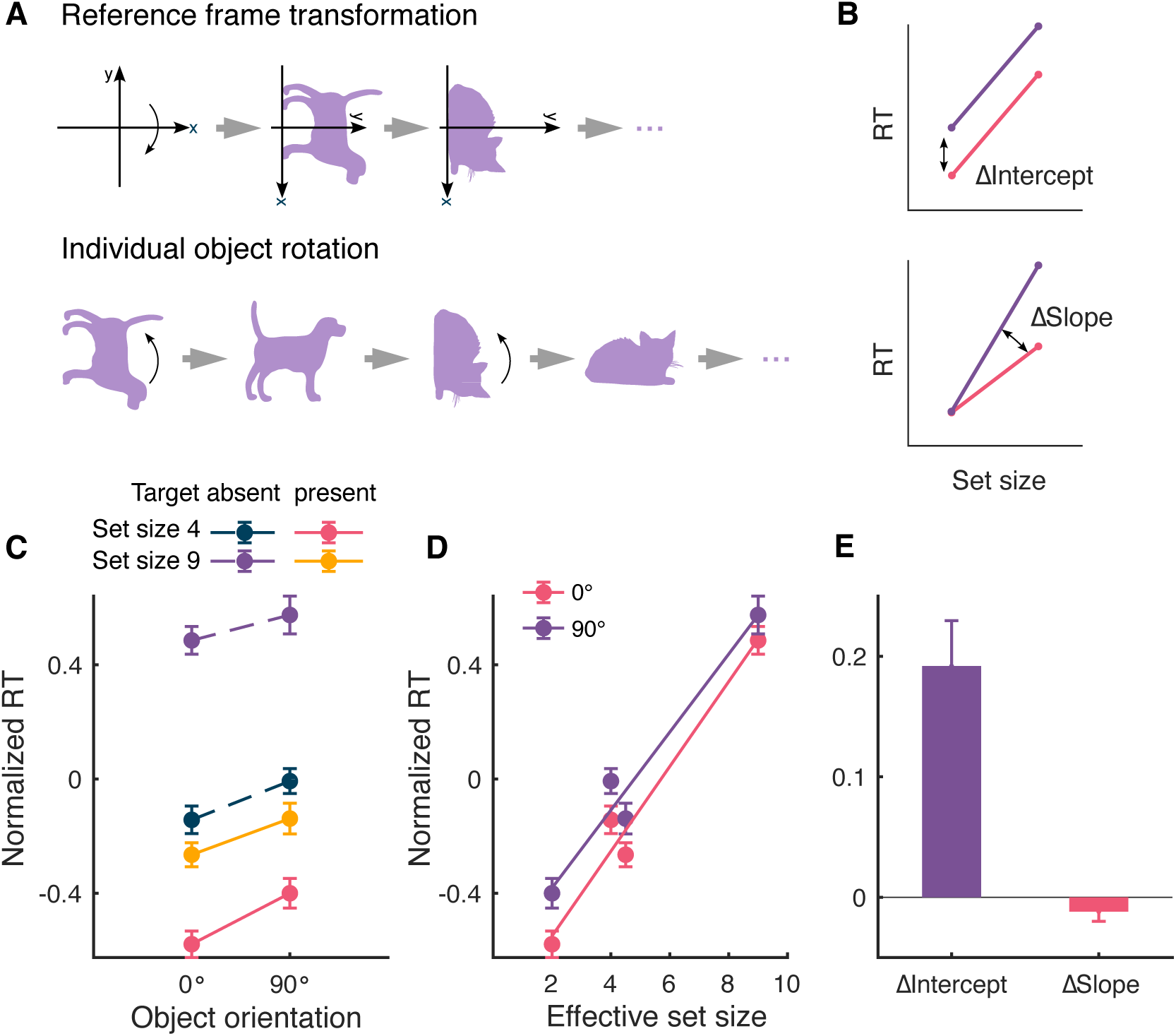
Illustrations of two possible mechanisms for orientation dependency and additional results for Experiment 1. A, Top row, an internal reference frame is rotated to align with objects at the beginning of each trial. Bottom row, each object is mentally rotated to a familiar orientation during serial search. B, Top, reference frame transformation predicts an orientation dependency on the intercept of the RT-set size relationship, as the transformation step is independent of the number of objects. Bottom, object rotation predicts a difference in slope between upright and tilted objects, because each object requires additional time for mental rotation. C, Longer normalized response time was found for both target-absent (dashed lines) and present trials (solid lines) for set sizes 4 (dashed blue and solid red lines) and 9 (dashed purple and solid yellow lines). D, Response time varied as a function of effective set size for upright (red) and tilted objects (purple). Lines represent linear fit to the data. E, Difference in intercept and slope of the response time–effective set size functions shown in D. Error bars indicate S.E.M.

Our linear mixed-effects model showed that the interaction between object orientation and set size was not significant (estimate=-1.30×10^−4^, 95%CI=[−3.39×10^−4^, 7.91×10^−5^], *t*(10029)=-1.22, *p*=0.223). Furthermore, we observed a significant difference in the intercepts of the RT–set size function for upright and horizontal objects (Figure 2D; *m*=-0.198, 95%CI=[−0.311, −0.086], *t*(19)=-3.68, *p*=0.00159, paired t-test), but not in the slopes (Figure 2D; *m*=0.009, 95%CI=[−0.008, 0.0261], *t*(19)=1.11, *p*=0.281). These findings suggest an internal reference frame transformation, rather than individual object rotation.

We also observed a significant difference between target-present and target-absent trials (Figure 3C; estimate=-0.160, 95%CI=[−0.291, −0.0283], *t*(10029)=-2.383, *p*=0.0172) and a significant interaction between target presence and set size (estimate=-0.0655, 95%CI=[−0.0846, −0.0464], *t*(10029)=-6.730, *p*=1.793×10^−11^). Interestingly, target presence did not significantly interact with object orientation (Figure 3C; estimate=3.85×10^−4^, 95%CI=[−1.71×10^−3^, 2.48×10^−3^], *t*(10029)=0.360, *p*=0.719). To reconcile these observations, we computed the effective set size to better characterize the relationship between set size and target presence. In target-absent trials, an observer must process all objects in the scene before asserting the target is absent. Therefore, the effective set size equals the actual set size. On the other hand, in target-present trials, the search can end as soon as the target is found. On average, only half of the objects would be processed^2^. As a result, the effective set size in target-present trials is half the actual set size. This 1:2 RT ratio between target-present and target-absent trials is well-documented in serial search^2,55^. We hypothesized that RT increases linearly with effective set size and that object orientation affects the intercept, but not the slope, of the RT– effective set size function.

Indeed, we found a strong correlation between effective set size and normalized RT (Figure 3D; *r*=0.871, *p*=8.95×10^−26^ for upright objects, *r*=0.816, *p*=3.22×10^−20^ for tilted objects, Pearson’s correlation). Importantly, the intercept, but not the slope, of the RT– effective set size function differs significantly between the two object orientations (Figure 3E; *m*=-0.192, 95%CI=[−0.271, −0.113], *t*(19)=-5.09, *p*=6.44×10^−5^ for intercepts; *m*=0.0119, 95%CI=[−0.00478, 0.0287], *t*(19)=1.49, *p*=0.151 for slopes, two-tailed paired t-test), consistent with our findings of the RT–set size function.

Together, these results demonstrate that the effects of object orientation on RT were independent of set size, which suggests a process separate from the core serial search. These findings are consistent with the hypothesis of reference frame transformation^25^, in which the observer aligns an internal reference frame with objects before starting the recognition or search processes.

### Allocentric cues contribute to orientation dependency

The findings in the previous section suggest that an internal reference frame is transformed to facilitate visual search for tilted objects. What cues might determine this internal reference frame during a real-world search? Two general types of reference frames might be involved: egocentric and allocentric reference frames^33,40,41^. In egocentric reference frames, objects are represented relative to the retina, head, or body. In allocentric reference frames, visual context and gravitational cues may be used as references^15,36,43,50,51,53,56,57^. To quantify the effects of different reference frames, we manipulated the visual context and egocentric cues by leveraging a virtual reality system consisting of a flight simulator and a head-mounted display (Figure 4).

**Figure 4.**
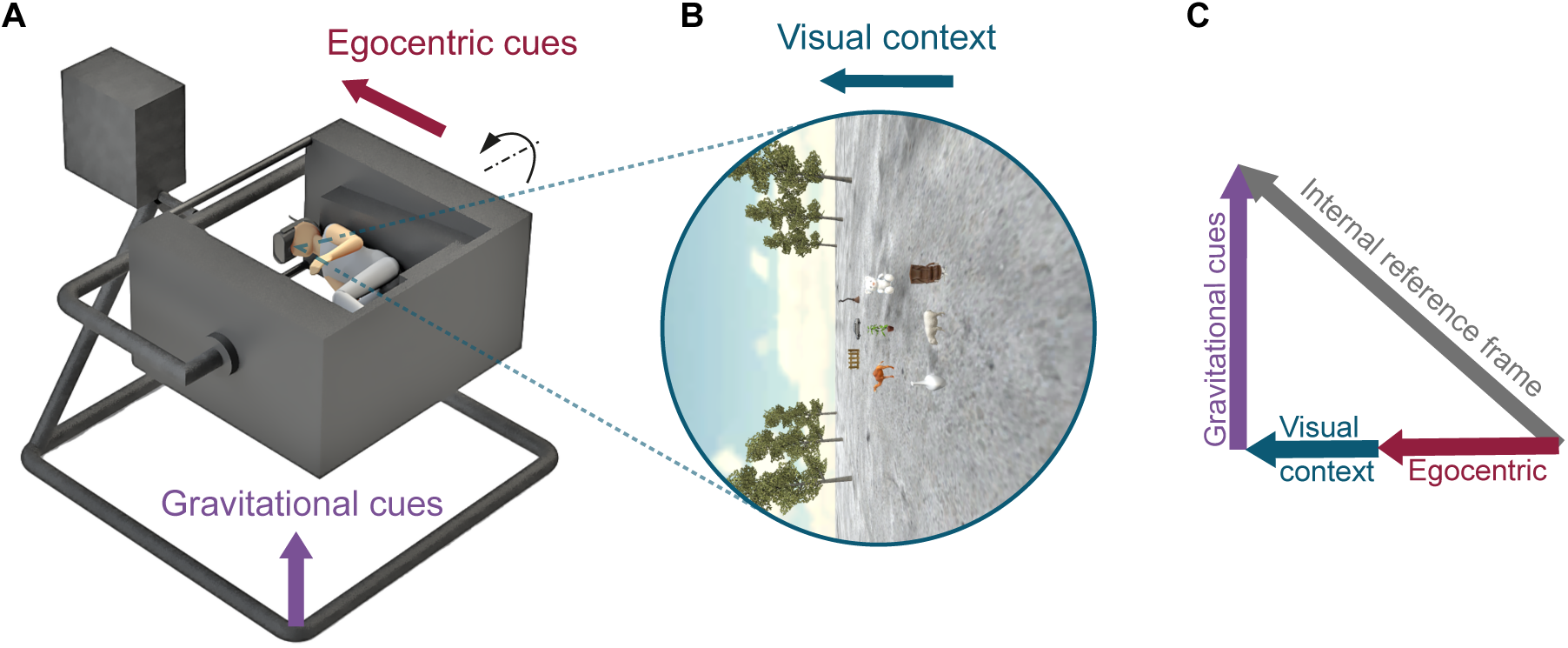
Apparatus in Experiment 2. A, Participants sat in a flight simulator, which allowed full body roll on a block-by-block basis, separating the egocentric reference cues from visual context and gravitational cues. B, The virtual scene presented on the head-mounted display was rotated independently to manipulate visual context. C, The internal reference frame can be modeled as a vector sum of visual context, egocentric, and gravitational cues with unknown weights. While not directly manipulated, gravitational cues can be dissociated from the other two cues by rotating the visual context and the participant’s body in the same direction.

In Experiment 2, four conditions were interleaved across blocks, including 1) a Baseline condition in which neither the visual context nor the flight simulator rotated, and all three reference cues were aligned (Figure 5A); 2) a Visual context condition in which the visual scene was rotated by 90° clockwise while the participants remained upright (Figure 5B); 3) an Egocentric condition in which the flight simulator rotated by 90° clockwise while the visual context remained upright relative to the world, such that egocentric cues differed from the other two cues (Figure 5C); and 4) a Gravitational condition, in which both the flight simulator and visual context rotated by 90° clockwise in the same direction, such that gravitational cues differed from these two cues (Figure 5D).

**Figure 5.**
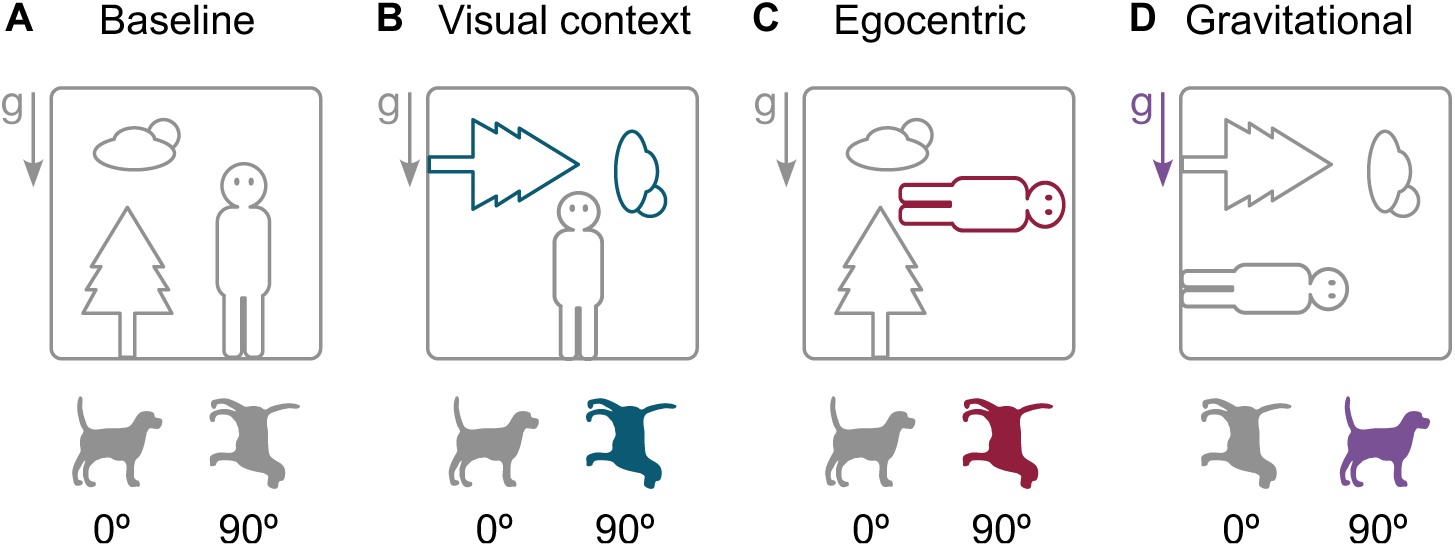
Experimental conditions in Experiment 2. A, Baseline condition, in which visual context (represented by the tree and cloud), egocentric (the observer’s head and body), and gravitational (gray arrow, g) cues were all upright. 0° object orientation indicated upright with respect to all three cues. B, Visual context condition, in which only the visual context was rotated by 90°. In this case, 90° became upright with respect to the visual context, and 0° was upright with respect to the other two cues. C, Egocentric condition, in which the observer’s full body was rotated by 90°. Here, 90° was upright with respect to egocentric cues, while 0° remained upright to the other two cues. D, Gravitational condition, in which both visual context and the observer’s body were rotated by 90° in the same direction. In this condition, 0° was upright to visual context and egocentric cues, and 90° was upright with respect to gravity.

Object orientation was defined relative to gravity in the Baseline, Visual context, and Egocentric conditions, and relative to the body in the Gravitational condition (Figure 5, bottom row). These coordinates were chosen such that any changes in orientation dependency can be attributed to the rotation of the reference cue under study in each condition. For example, if RT is shorter for upright than tilted objects (with respect to gravity) in the Visual context condition, it would suggest that visual context has no contribution to the orientation dependency. In contrast, if similar RT is found between upright and tilted objects, it would indicate that visual context has a significant effect that cancels out the orientation advantage. Finally, if tilted objects, which would appear upright relative to the visual context, have shorter RT in this condition, it would suggest an even stronger contribution of visual context that reverses the orientation dependency.

We found a significant main effect of object orientation on normalized RT (Figure 6A; estimate=1.64×10^−3^, 95%CI=[8.13×10^−4^, 2.47×10^−3^], *t*(9865)=3.89, *p*=1.02×10^−4^, linear mixed-effects model; see Methods). Importantly, there were significant interactions between object orientation and visual context (estimate=-3.11×10^−5^, 95%CI=[−4.40×10^−5^, −1.81×10^−5^], *t*(9865)=-4.69, *p*=2.73×10^−6^), between object orientation and egocentric cues (estimate=-2.25×10^−5^, 95%CI=[−3.54×10^−5^, −9.50×10^−6^], *t*(9865)=-3.40, *p*=6.84×10^−4^), and among object orientation, visual context, and egocentric cues (estimate=6.61×10^−5^, 95%CI=[4.58×10^−7^, 8.65×10^−7^], *t*(9865)=6.36, *p*=2.14×10^−10^). These findings suggest that all three reference cues influenced the orientation dependency observed in our visual search task.

**Figure 6.**
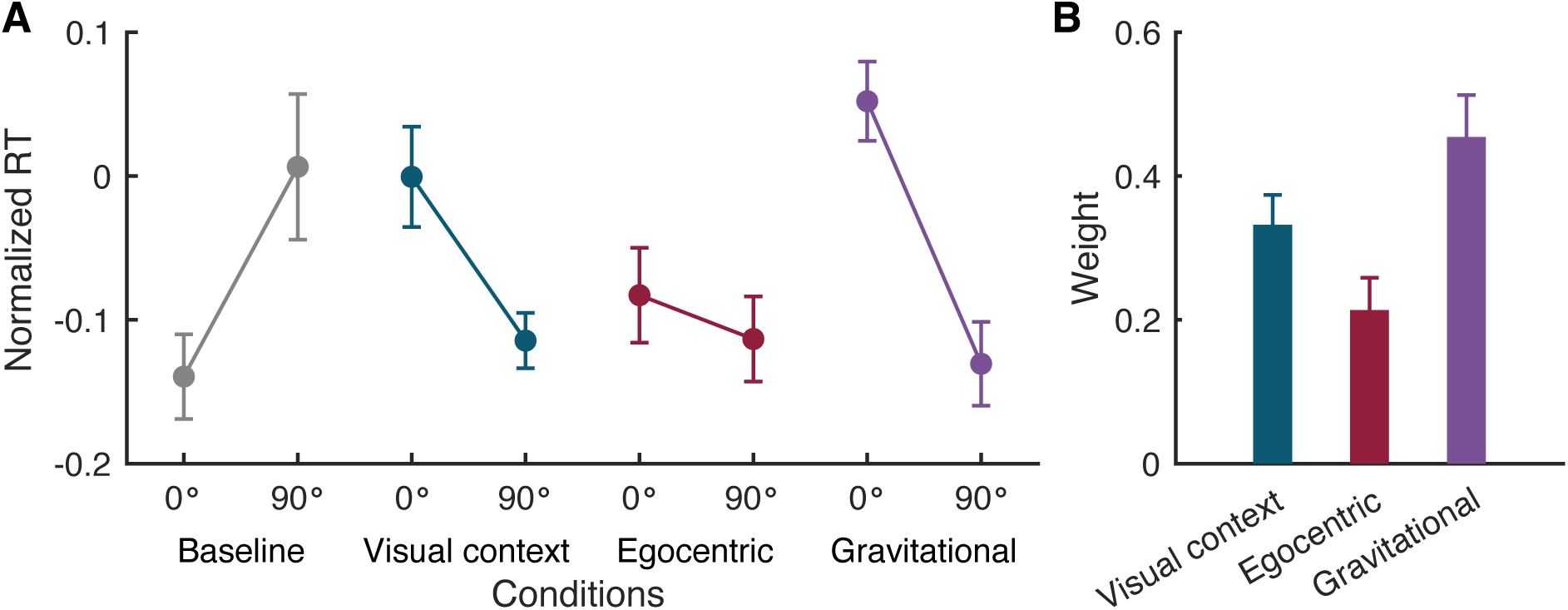
Results for Experiment 2. A, Normalized response time in each experimental condition. 0° and 90° represent upright and tilted objects, respectively. In each condition, object orientation was defined relative to the reference cues not under experimental manipulation (i.e., gravitational reference frame in the Baseline, Visual context, and Egocentric conditions; egocentric reference frame in the Gravitational condition; see Figure 5). B, Weights for each reference frame inferred from changes in orientation dependency. Error bars represent S.E.M.

In the Baseline condition, the effect of object orientation is consistent with that in Experiment 1, with faster RT for upright than tilted objects (Figure 6A, gray; *t*(9)=-3.6705, *p*=0.0052, 95%CI=[−0.2358, −0.0560], two-tailed paired t-test). In the Visual context condition, changing the orientation of visual context completely reversed the orientation dependency, resulting in a faster response to objects that were tilted with respect to egocentric and gravitational reference frames (Figure 6A, dark blue; *t*(9)=3.2334, *p*=0.0103, 95%CI=[0.0342, 0.1936], two-tailed paired t-test). This indicates that visual context can largely overwrite the orientation defined by the other two cues. In contrast, rotating the egocentric cues resulted in similar RTs between the two object orientations (Figure 6A, dark red; *t*(9)=1.0519, *p*=0.3203, 95%CI=[−0.0351, 0.0960], two-tailed paired t-test). This result suggests that egocentric cues were sufficient to eliminate orientation dependency, but not strong enough to reverse it. Finally, orientation dependency was also reversed in the Gravitational condition (Figure 6A, purple; *t*(9)=5.6931, *p*=0.0003, 95%CI=[0.1100, 0.2551], two-tailed paired t-test), suggesting a strong effect of gravitational cues in determining the effect of object orientation in visual search.

We computed the relative weights for these three reference frames by comparing an orientation dependency index for each condition (Figure 4C and 6B; see Methods). The weights for all three references were significantly higher than zero (estimate=0.33, 95%CI=[0.23, 0.43], *t*(27)=7.17, *p*=1.04×10^−7^ for visual context; estimate=0.21, 95%CI=[0.12, 0.31], *t*(27)=4.61, *p*=8.77×10^−5^ for egocentric cues; estimate=0.45, 95%CI=[0.36, 0.55], *t*(27)=9.80, *p*=2.18×10^−10^ for gravitational cues; linear mixed-effects model). An analysis of variance revealed that the weights varied significantly across reference cues: *F*(3, 27)=56.22, *p*=9.71×10^−12^. Gravitational cues had the highest weight, visual context ranked second, and the weight for egocentric cues was the lowest (Figure 6B). Tukey’s HSD test revealed that the weight for gravitational cues is significantly higher than that for egocentric cues (mean difference=0.24, 95%CI=[0.07, 0.41], *p*=0.0047), but the differences between other pairs of weights were not significant (mean difference=0.12, 95%CI=[−0.05, 0.29], *p*=0.217 between visual context and egocentric cues; mean difference=0.12, 95%CI=[−0.05, 0.29], *p*=0.200 between gravitational and visual context cues). It is worth noting that these were relative weightings across three types of reference cues, and although ranked the lowest, egocentric cues had considerable effects, as shown by the absence of orientation dependency in the Egocentric condition (Figure 6A, compare gray and dark red).

These results demonstrated a drastic change in orientation dependency when different reference frames were manipulated. This confirms our findings in Experiment 1 that an internal reference frame is involved during visual search for tilted objects. Furthermore, these results suggest that multiple ego- and allocentric reference cues constitute the internal reference frame, which may help to establish a stable representation of the world (see Discussion).

## Discussion

In this study, we investigated how object orientation and reference cues influence visual search performance in natural scenes. Our results revealed that participants were more efficient at locating upright objects compared to tilted ones, demonstrating an orientation dependency in real-world contexts. We considered two possible mechanisms underlying this orientation dependency: internal reference frame transformation and individual object rotation^25^. Further analysis of the relationship between RT and set size revealed that only the intercept, but not the slope, of the RT–set size function differed between upright and tilted objects. This finding rules out the hypothesis of individual object rotation, which predicts a change in the slope of the RT– set size relationship. Instead, our findings suggest that the brain employs an internal reference frame for visual processing, and this internal reference frame is rotated to match the orientation of objects before the search.

By rotating the participants’ bodies and the visual scene in virtual reality, we showed that multiple reference cues, including egocentric, visual context, and gravitational, contribute to the internal reference frame. Notably, allocentric reference frames, namely visual context and gravitational cues, had greater influences on the orientation dependency than egocentric cues. This suggests that multisensory integration and higher-order cognitive mechanisms play a critical role in establishing the internal reference frame during visual search in complex, real-world environments.

What is the advantage of using an internal reference frame? Transforming an internal reference frame, rather than sequentially rotating each object, significantly reduces the computational complexity of visual search in the real world. By realigning the reference frame once before initiating a search, the process requires a fixed amount of cognitive resources that do not depend on set size. This allows the brain to process complex natural scenes in a more energy-efficient manner. In contrast, if objects are rotated individually, each additional object would necessitate additional cognitive resources, making the process more resource-intensive as the visual scene becomes more complex. This study, consistent with prior studies on reference frames^17,24,33–38,41,42,50,57^, demonstrates that the nervous system adapts to environmental statistics and efficiently processes information based on task demands. Our findings suggest that internal reference frame transformation might be another strategy the brain uses to improve visual search efficiency in real-world environments, complementing existing factors such as top-down attention, stimulus saliency, prior knowledge, and semantic context^5,6,9–13,56,58^.

Visual information is primarily encoded in a retinotopic space in the early stages of visual processing^59–61^. However, a retinotopic representation is often unstable due to movements of the eyes, head, and body as we navigate the environment. To form a stable representation of the world, information about posture and self-motion—including efference copy of motor commands and sensory inputs from multiple modalities—is integrated to transform visual inputs into allocentric coordinates^36,38,39,42,43,50,51,53,57,62–67^. This transformation from egocentric to allocentric reference frames might be initiated in V1^68–70^, continued along both the dorsal visual pathway to the parietal cortex^31,34,39,40,42–45,49,62,71–74^, and the ventral pathway to the inferotemporal cortex^46,47,75^.

Our results show that real-world visual search relies heavily on gravitational cues, which may include vestibular signals and proprioception^43,51,57^. These findings are supported by a recent neurophysiological study showing that objects are predominantly encoded in a gravitational reference frame in the primate inferotemporal cortex, an area central to object processing^46^. In addition, neurons in the anterior inferotemporal cortex exhibit tuning for visual gravity cues akin to the visual context in our study^47,75^. Gravity orientation tuning has also been found in the anterior thalamus, cerebellum, and other regions in primates and rodents^76–78^. Together, these studies provide strong neural evidence for an internal reference frame that integrates visual and nonvisual cues.

Consistent with previous studies that demonstrated the influence of gravity on visual perception^36,48,51,79^, we show for the first time that gravitational cues can outweigh egocentric cues in visual search. The greater reliance on these cues might be driven by the complexity of natural objects and scenes, which requires the engagement of high-level visual areas^9,46,47,75^. The visual system might rely more on retinotopic or egocentric reference frames when processing simple visual features such as oriented bars and intersecting line segments^14,16,24,41,79^. An interesting future direction is to examine the relative weightings of different reference cues in tasks involving stimuli ranging from simple to complex^48^. It is possible that the nervous system stores information in multiple reference frames and adaptively chooses an optimal reference based on the context^34,44,45,49^.

Another possible explanation for the higher weighting on gravity is the involvement of an internal reference frame in our task. Previous studies primarily used tasks that involved only a single stimulus on the display^36,79^. In these tasks, a reference frame transformation may not be necessary, as mental rotation of a single object would be equally efficient. It is therefore natural that gravity influences perception less when these reference frames are not explicitly used. In contrast, our visual search promotes the use of reference cues rather than mental rotation of each object, leading to greater engagement of other sensory systems needed for a robust internal reference frame. However, as discussed above, gravity-centered representation might be a universal property of the higher visual areas, regardless of the task context^46,47,75^. Future work that examines the involvement of an internal reference frame in various contexts would provide valuable insights.

Due to technical limitations, we did not measure eye positions during the experiments. Roll vestibular-ocular reflex produces torsional eye movements in the direction opposite to body rotation^80^, which may reduce the rotation of retinotopic coordinates in the Egocentric and Gravitational conditions in Experiment 2. This reduction in egocentric orientation might explain the smaller weights for egocentric reference cues. However, because enough time was given for participants to adapt to new body postures after each body rotation, we believe that only static torsion, not torsional eye movement, was involved during task performance. The amount of static torsion is 10°-12° maximum^80^, which could account for only ∼11% of errors in our estimation of weights. It has also been shown that static tilted visual images could not induce torsional eye movement^80^. Thus, our results in the Visual context condition were unlikely to be influenced by torsion. Future work using reliable eye-tracking in virtual reality would further our understanding of how eye movements interact with the rotation of multiple reference cues.

In summary, this study demonstrates that humans use an internal reference frame in real-world visual search. Using a virtual reality system, we show that multiple allocentric and egocentric reference cues shape this internal reference frame, providing a novel explanation for visual search efficiency in naturalistic environments. Our findings suggest that real-world visual search offers valuable insights into high-level cognitive processes that involve multiple sensory modalities.

## Methods

### Participants

Twenty participants (seven males and thirteen females, mean age 20.40, SD=0.49) with normal or corrected-to-normal vision participated in Experiment 1, and ten of them (five males and five females, mean age 20.90, SD=0.80) participated in Experiment 2. All subjects were new to psychophysical experiments and were unaware of the purpose of the study. Informed written consent was obtained from each participant before data collection. The study was approved by the Institutional Review Board at East China Normal University.

### Apparatus

A head-mounted display (HTC Vive, first generation, HTC and Valve Corporation) was used for stimulus presentation. The display had a mean luminance of 61.15 cd/m^2^, a resolution of 1080×1200 pixels, a refresh rate of 90 Hz, and a pixel size of ∼0.09°×0.09°. The field of view of the head-mounted display was approximately 111.5°. Visual stimuli were generated in Unity3D (Unity Technologies, version 5.6.2). Participants sat on a chair inside a custom flight simulator (Zhuoyuan Co.) and were secured by a seatbelt and additional padding around the head and body (Figure 4A). A wireless joystick was held by the participants to respond to the task. The flight simulator could be rotated along the roll axis at low speed.

### Stimuli

The visual stimuli consisted of an outdoor scene with objects placed on the ground (Figure 1). A matrix of either four or nine objects was presented at the center of the screen in an area measuring ∼49°×53°. The size of each object was around 4.8°×4.8° on the display and was the same for both set sizes, 4 and 9. The spacing between adjacent objects was the same for both set sizes, such that differential effects of visual crowding could be eliminated. Objects included large animals and furniture with a canonical orientation. Their orientation was either 0° (upright) or 90° (horizontal), with all objects sharing the same orientation during each trial. Objects presented on each trial were randomly selected from 89 categories, each comprising two variations with different shapes and colors. In target-present trials, the search target was randomly selected from the presented objects; in target-absent trials, the target was randomly selected from the absent objects. All 3D models used in the experiments were acquired from the Unity Asset Store and were scaled to approximately the same 3D volume.

### Experimental conditions

In Experiment 1, two set sizes, 4 and 9, were interleaved across blocks. Each block consisted of 72 trials. Two object orientations, horizontal and upright, were randomly intermixed and counter-balanced across trials. In half of the trials, the array of objects did not include the cued target (target-absent trials). Target-present and target-absent trials were counter-balanced and randomly intermixed within each block. The flight simulator and visual context did not rotate throughout the experiment. Each participant completed two sessions with four blocks in each session.

In Experiment 2, four reference frame conditions were introduced (Figure 5). 1) In the Baseline condition, the flight simulator and visual context remained upright relative to gravity (Figure 5A). 2) In the Visual context condition, the visual scene was rotated 90° in either direction relative to gravity (Figure 5B). 3) In the Egocentric condition, the flight simulator rotated 90° while the visual context remained upright (Figure 5C). 4) In the Gravitational condition, both the flight simulator and visual context rotated 90° in the same direction, such that only gravitational cues differed from the other two cues (Figure 5D). These four conditions were separated into blocks and randomly interleaved. The set size was 9 throughout the experiment. Object orientation and target presence conditions were interleaved in the same way as in Experiment 1. Each participant completed two sessions with eight blocks in each session. Importantly, the head-mounted display covered the entire field of vision and did not move relative to the observer’s head throughout the experiment, allowing control over the visual information received by the participants regardless of their body orientation. Note that body rotation only occurred between each block of trials, and we allowed participants to adapt to the new body orientation before resuming the experiment. Therefore, we expect the eyes to reach a steady state that involves only a small amount of torsion, if any^80^.

### Procedure

On each trial, a word cue was presented for 500 milliseconds, indicating the search target for the current trial (Figure 1). After a variable inter-stimulus interval (ISI), a set of objects was presented on the screen. The ISI was randomly selected from a set of discrete values ranging from 500 to 1000 milliseconds with a step of 100 milliseconds. Participants were asked to report whether the target was present by pressing one of the two buttons on the joystick. The objects disappeared after a response was made. If a response was not made within 1500 milliseconds, the objects would disappear, and participants would still need to respond before proceeding to the next trial. This 1500-ms stimulus duration encouraged participants to respond as fast and accurately as possible, which possibly prevented them from revisiting the same object multiple times. On average, most correct responses were made within the stimulus presentation period (mean = 1094±375 ms for Experiment 1; mean = 1208±433 ms for Experiment 2). The next trial started 1000 milliseconds after a response was made.

### Data preprocessing

For each participant, trials with RT greater than 4 median absolute deviations (MADs) from the median were removed from further analysis. MAD was used because it is more robust to outliers than standard deviation. In addition, trials with RT shorter than 50 ms were classified as random guesses and were therefore removed from analysis. On average, 2.67% of the trials were removed. We then normalized RTs by taking the Z-score for each participant and only used RTs from correct trials for further analyses.

### Data analysis

#### Discriminability

The discriminability of each subject was calculated as: *d*′ = *Z*(*hit*) − *Z*(*false alarm*), where *Z*(*hit*) was the Z score of the hit rate and *Z*(*false alarm*) was the Z score of the false alarm rate.

#### Effective set size

In target-absent trials, an observer must identify all objects before responding. Therefore, the effective set size would be the same as the actual set size. In contrast, when the target is present, only half of the objects, on average, need to be processed before the target is identified. Therefore, the effective set size would be half of the actual set size in target-present trials^2^.

#### RT–set size function

A linear function, *z*(*RT*) = *b*_1_ + *b*_2_*x*, was fit to the relationship between (effective) set size *x* and the z score of RT, where *b*_1_ and *b*_2_ are the intercept and slope of the function, respectively. Regression coefficients were computed using ordinary least squares in MATLAB (MathWorks, MA).

#### Linear mixed-effects models

Because of the nature of RT data, there is a large variability across individuals and trials^81^. We modeled RT using linear mixed-effects models that considered participants and object categories as random effects on the RT intercept and experimental conditions as fixed effects. These models were fit to trial-level data using the *fitlme* function in MATLAB (MathWorks, MA).

For Experiment 1, we modeled normalized RTs for each trial using three variables and all permutations of interactions between them as fixed effects, including target presence, object orientation, and set size. Random effects on intercept included participant identity and object category in each trial. The object orientation factor in the model was considered a measure of orientation dependency. The interaction between object orientation and set size was considered the effect of orientation on the slope of the RT-set size function. The three-way interaction between object orientation, set size, and target presence was considered the effect of orientation on the slope of the RT-effective set size function.

For Experiment 2, we modeled normalized RTs using object orientation and the orientations of visual context and egocentric reference frame as fixed effects, and the random effects were the same as in Experiment 1. Interactions between Object orientation x Visual context and Object orientation x Egocentric were considered contributions of visual context and egocentric cues, respectively. The three-way interaction between Object orientation x Visual context x Egocentric was considered the effect of gravitational cues.

#### Contributions of reference frames

To quantify the relative weights of each reference frame in Experiment 2, we first computed an orientation dependency index (ODI) as the difference in z-scored RT for horizontal and upright objects:

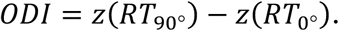

Weights for reference frames were then computed as the change in ODI in each condition compared to the baseline:

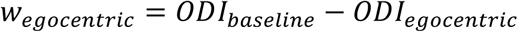

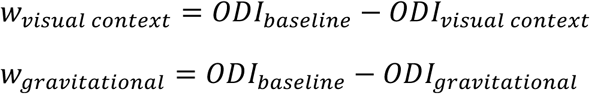

Finally, the weights were normalized by their sum for each participant. Note that object orientation was defined relative to the reference frame that was not under manipulation in each experimental condition (i.e., gravitational reference in Baseline, Visual context, and Egocentric conditions; egocentric reference in the Gravitational condition; see Figure 5). A linear mixed-effects model was fit to the weights with participant identity as a random effect: Weight ∼ Visual context + Egocentric + Gravitational + (1|Subject ID). This provides a statistical assessment of the significance of each weight. An analysis of variance on the model was used to measure variability across reference cues.

## Acknowledgments

We (Y. C. and Z. X.) performed the experiments while we were students in Dr. Shuguang Kuai’s laboratory at East China Normal University. We thank Dr. Kuai for his support (including the National Natural Science Foundation of China grants 31771209 and 3151160 to S. K.) and Dr. Greg DeAngelis for helpful comments on the manuscript.

## Data and code availability

All data and code used in this study are available at: https://osf.io/zmt2g/?view_only=ac02a214e86846eead93729b3dc70e6d.

